# Variable prevalence of protective *Spiroplasma* infection over time in two natural populations of *Drosophila hydei*

**DOI:** 10.1101/2024.07.31.606006

**Authors:** Jordan E. Jones, Rebecca Court, Daisuke Kageyama, Darren J. Obbard, Gregory D. D. Hurst

**Affiliations:** Institute of Infection, Veterinary and Ecological Sciences, University of Liverpool, Liverpool, L69 7ZB, UK; National Agriculture and Food Research Organization (NARO), 1-2 Owashi, Tsukuba, Ibaraki 305-0851, Japan; Institute of Ecology and Evolution, University of Edinburgh, Ashworth Laboratories, Charlotte Auerbach Road, Edinburgh, EH9 3FL, UK

**Keywords:** Spiroplasma, Drosophila hydei, Infection prevalence, Protective symbiosis

## Abstract

The temporal dynamics of protective symbionts have rarely been characterized outside of aphid hosts. Here, we determine the prevalence of *Spiroplasma* in two populations of *D. hydei* where *Spiroplasma* infection had been previously recorded (UK and Japan). We observe that infection in both populations is variable over time and confirm the persistence of *Spiroplasma* in the UK population for 9 years. Thus, variable prevalence over time appears to be a common feature of these symbioses.

## Introduction

Many insects harbour heritable symbiotic bacteria that can provide a source of rapid and dynamic adaptation in their hosts. Recording of longitudinal prevalence data can provide insight into how symbiont-mediated traits shape infection and capture evolutionary events unfolding within relatively short timescales. For example, *Rickettsia* increased in south-western populations of the whitefly, *Bemisia tabaci*, from 1% to 97% in just 6 years due to infected individuals developing faster and producing more offspring than their uninfected comparators (Himler *et al*., 2011). Similarly, the prevalence of *Spiroplasma* in *Drosophila neotestacea* was observed to rapidly spread from the east to the west coast of the USA driven by its ability to protect hosts against the harmful effects of pathogenic nematodes (Jaenike *et al*., 2010; Cockburn *et al*., 2013). Symbiont frequencies can also fluctuate rapidly over shorter time scales, as demonstrated by highly variable symbiont frequencies observed across 6 months in wild aphid populations in the USA (Smith *et al*., 2015).

*Spiroplasma* is a maternally inherited bacteria of the fruit fly, *Drosophila*. Although initially known for its ability to kill males (Montenegro *et al*., 2005), *Spiroplasma* is now also considered a beneficial partner of *Drosophila*, protecting infected individuals against pathogenic nematodes and parasitic wasps through a combination of toxin secretion and resource competition for lipids (Paredes *et al*., 2016; Ballinger and Perlman, 2017). Unlike in some *Drosophila* species, *Spiroplasma* does not cause male-killing in Drosophila hydei, but can protect against Leptopilina parasitoid wasps, killing the developing wasp and rescuing the host (Xie *et al*., 2010). Longitudinal data of *Spiroplasma* frequency in *D. hydei* is limited to a single study which found infection frequency to remain stable at 15% over two consecutive years in a UK population (Corbin *et al*., 2021). Geographical prevalence data is more commonly described and has shown that *Spiroplasma* in *D. hydei* can reach intermediate frequencies in natural populations. For example, *Spiroplasma* frequency ranged from 23-66% in Japanese populations (Kageyama *et al*., 2006) and similar infection frequencies were observed in populations from western USA ranging from 26-60% (Watts et al., 2009). *Drosophila hydei* is known to be infected with two strains of *Spiroplasma* known as *sHy1* and *sHy2*, although *sHy2* infection is less commonly observed and records to date are limited to North America (Mateos et al., 2006).

In this study, we determine the infection prevalence of *Spiroplasma* from two populations of *D. hydei* from which infection frequency has previously been recorded; i) Motts Mill, UK and ii) Tsukuba, Japan. *Spiroplasma* infection prevalence was found to be 15% during the summers of 2014 and 2015 in the UK population (Corbin *et al*., 2021), whereas a higher infection prevalence of 26% was recorded in the Japanese population in 2005 (Kageyama *et al*., 2006). To this end, we sampled *D. hydei* from Motts Mill, UK (collected in 2021 and 2023) and Tsukuba, Japan (collected in 2021), and tested them for Spiroplasma infection using PCR assays to i) determine whether *Spiroplasma* infection continues to persist in these populations since the previous sampling year and, ii) determine whether *Spiroplasma* infection frequency remains stable or is dynamic across sampling years.

## Materials and methods

### Drosophila hydei collection from field

*Drosophila hydei* samples from the UK (Motts Mill, 51.1 N, 0.164 E) were collected in July and September 2021 (561 individuals), and during August 2023 (230 individuals). Samples collected in 2021 were stored in ethanol at -20°C and samples collected in 2023 were stored dry at -20°C. *Drosophila hydei* samples from Japan (Tsukuba, 36.03 N, 140.09 E) were collected between July and September 2021 (156 individuals). Samples were stored dry at -80°C. All samples were sexed and screened for the presence of ectoparasitic mites, which were removed where found.

### DNA extraction and PCR for Spiroplasma infection

DNA from whole fly bodies were extracted using Promega Wizard® genomic DNA kit following the manufactures protocol for ¼ of the measures given for animal tissue. DNA template quality was tested for each sample using *CO1* amplification using the primers *‘HCO’* and *‘LCO’* (Table 1). Following this, all samples were screened for *Spiroplasma* infection using the primers *‘SpoulF’* and *‘SpoulR’* (Table 1). Samples which did not amplify for the *CO1* gene were excluded as of poor quality (UK 2023: 1 sample; UK 2021: 2 samples; Japan 2021: 5 samples).

**Table 1:**
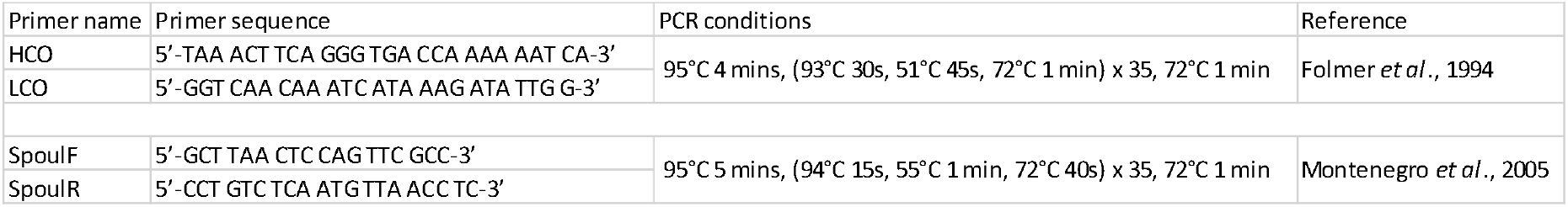
Primer sequences and PCR cycling conditions used in the study.

### Sanger sequencing for Spiroplasma strain confirmation

To determine the strain of *Spiroplasma* found in the samples collected, a subset of amplicons for *Spiroplasma*-positive samples were sent for sequencing (UK 2021: 14 samples; UK 2023: 18 samples; Japan 2021: 4 samples). To this end, PCR products underwent an ExoSAP digest clean up to remove excess primers, and 5 μl of PCR product was added to a mixture containing 0.2 μl Shrimp alkaline phosphate, 0.05 μl of Exonuclease I, 0.7 μl 10X RX Buffer and 1.05 μl of molecular grade water. Samples were then incubated for 45 min at 37 °C followed by 15 min at 80 °C and sent for Sanger sequencing using the SpoulF primer. *‘sHy1’* and *‘sHy2’* strains were distinguished through the presence of two SNPs at positions 34 and 49 respectively on the 16S rRNA amplicon (*sHy1*: base 34 of amplicon = cytosine and 49 = guanine; *sHy2*: base 34 of amplicon = thymine and 49 = adenine).

### Statistical analysis

All statistical analyses were conducted using the statistical software, R version 4.0.2 (R Core Team, 2022). Using *Spiroplasma* prevalence data from previous studies (Corbin *et al*., 2021; Kageyama *et al*., 2006) and equivalent data from the same populations collected in this study, we determine whether *Spiroplasma* infection significantly differed by year of collection. A generalised linear model with binomial errors was used to determine the effect of year of collection and sex on *Spiroplasma* infection prevalence of wild caught *D. hydei* from the UK population. The *“Anova”* function in the *“car”* package version 3.1–0 with *F* tests were used to assess significance (Fox and Weisberg, 2019). Due to the absence of male infection data in the comparable Japan data set, a Fisher’s exact test (2 × 2 contingency table, year of collection x infection status) was performed to determine whether there was a significant difference between *Spiroplasma* infection frequency and year of collection of *D. hydei*.

## Results

In the UK population, year of collection had a significant effect on *Spiroplasma* infection status of *D. hydei* (Binomial GLM: F = 12.8, df = 3, p = 0.033). There was no significant effect of *D. hydei* sex on *Spiroplasma* infection (Binomial GLM: F = 3.61, df = 1, p = 0.15). In the Japan population, there was a significant difference between *Spiroplasma* infection and year of collection of *D. hydei* (Fisher’s exact: p < 0.001). All samples were confirmed to be *‘sHy1’* indicating that fluctuations observed were in the frequency of a particular symbiont, not through a change in symbiont strains present.

**Table 2:** *Spiroplasma* infection prevalence data from wild caught *Drosophila hydei* from the UK and Japan.

| Population | Year | Male |  | Female |  | Total |  |  | Reference |
| --- | --- | --- | --- | --- | --- | --- | --- | --- | --- |
|  |  | *N* collected | *N* Infected | *N* collected | *N* Infected | *N* collected | *N* Infected | % Infected |  |
| Motts Mill, UK | 2023 | 146 | 46 | 83 | 21 | 229 | 67 | 29 | *This study* |
| Motts Mill, UK | 2021 | 340 | 67 | 219 | 41 | 559 | 110 | 20 | *This study* |
| Motts Mill, UK | 2015 | 93 | 15 | 90 | 12 | 183 | 27 | 15 | Corbin et al., 2021 |
| Motts Mill, UK | 2014 | 30 | 6 | 30 | 3 | 60 | 9 | 15 | Corbin et al., 2021 |
| Tsukuba, Japan | 2021 | 110 | 7 | 41 | 3 | 151 | 10 | 7 | *This study* |
| Tsukuba, Japan | 2005 | N/A | N/A | 108 | 28 | 108 | 28 | 26 | Kageyama et al., 2006 |

## Discussion

There was a significant difference in the frequency of *Spiroplasma* infection between collection years of *Drosophila hydei* for both populations located in the UK and Japan, suggesting that *Spiroplasma* infection prevalence in *D. hydei* is dynamic across time. This study also confirms the persistence of *Spiroplasma* infection within a single UK *D. hydei* populations for 9 years.

Indeed, symbiont populations can undergo fluctuations in frequencies due to bottlenecks and resulting drift processes. Infection prevalence may also fluctuate due to selection pressures imposed by the environment. *Spiroplasma* is a protective symbiont that can enable survival of *D. hydei* following attack by *Leptopilina* parasitoid wasps. Higher attack rates by parasitoids in an area will increase the risk of parasitism to *Drosophila* and in turn, the prevalence of a protective symbiont would be expected to increase in response to selection imposed by the parasitoids. Indeed, this is observed within controlled experiments where *Spiroplasma* frequency increases within a population under constant wasp pressure, ultimately leading to fixation of the symbiont in highly attacked populations (Xie *et al*., 2015). However, for *Hamiltonella defensa* in wild aphid populations, longitudinal field studies reported that parasitism risk was not a key explanatory factor explaining the prevalence of the protective symbiont, (Smith *et al*., 2021; Gimmi *et al*., 2023). Whether the same is true for natural *Drosophila* populations remains to be determined.

The thermal environment is another factor likely to influence symbiont infection prevalence, due to symbiont infection sensitivity to temperatures. For example, *Spiroplasma* is temperature sensitive, becoming lost in populations of *D. hydei* passaged at below 18°C (Corbin *et al*., 2021). Although the mechanistic basis of thermal sensitivity is unclear, evidence from *Spiroplasma* in *D. melanogaster* suggests that *Spiroplasma* titre is reduced in flies reared at cooler temperatures, which can impact onward transmission, resulting in segregation and loss of infection (Jones and Hurst, 2023). Temperature and symbiont infection frequency data from natural populations is limited in *Drosophila*. However, the infection prevalence of the protective symbiont, *Hamiltonella defensa*, in natural populations of the black bean aphid was found to be best explained by the number of heat days that previous aphid generations had been exposed to (Smith *et al*., 2021; Gimmi *et al*., 2023), indicating that the thermal environment may represent an important factor underpinning yearly symbiont variation prevalence.

In this study we have confirmed the persistence of *Spiroplasma* infection within a single UK population across 9 years. Given the thermal sensitivity of *Spiroplasma* to temperatures below 18°C, which are often experienced in the UK, this continuity requires explanation. Little is known about the ecology of *D. hydei* within the UK. Occurrence data seems to suggest that *D. hydei* are observed all year round in the UK (Biological Records Centre, 2024). However, a caveat to this data is that not all occurrences supplied information on whether the *D. hydei* were observed outside or inside. Phenological data from *D. hydei* in Kansas, USA reports the early appearance of the species in April and a late disappearance in October (Gleason *et al*., 2019). Where the species resides over the cooler winter months is unknown but it is thought that human buildings or compost heaps may offer a winter refuge for the species (Spencer, 1941). Indeed, it is not well understood how well individuals with or without *Spiroplasma* infection survive over winter. The costs associated with harbouring infection may decrease their chances of survival through winter months. Furthermore, particularly harsh winters may also cause extreme bottlenecks in the symbiont population, further contributing to lower infection rates at the start of the spring and a dynamic prevalence over years.

## Acknowledgments

We would like to thank K and S Obbard for permission to collect specimens from Cherry Gardens Farm. This project was supported by funding from a NERC grant award (Grant number: NE/V011979/1) and a Globalink placement NERC award (Grant number: NE/V009834/1).

